# Network-level selection constrains the adaptive evolution of gene expression noise in regulatory genes in changing environments

**DOI:** 10.1101/2023.12.04.569843

**Authors:** Nataša Puzović, Julien Y. Dutheil

## Abstract

Gene expression levels vary from cell to cell, even between clonal cells in identical environmental conditions. This variance is termed “expression noise” and is partially genetically encoded and subject to selection. While gene expression noise is detrimental in constant environments where the expression level is under stabilizing selection, it can be beneficial in changing environments, when the phenotype is far from the optimum. However, expression noise propagates along the gene regulatory network, making the evolution of connected genes interdependent. Here, we explore how the position of the gene in the gene network constrains the evolution of its expression noise and expression mean under selection using in silico evolution experiments. We simulate the evolution of populations of model gene regulatory networks under directional and fluctuating selection on the gene expression level while allowing the basal expression level and expression noise level to mutate. We find that expression noise is only transiently favoured under directional selection, indicating a co-evolutionary dynamic between expression mean and noise during adaptation. On the other hand, high levels of noise are persistently maintained under fluctuating selection. Furthermore, target genes, regulated by other genes, were more likely to respond to selection and increase their gene-specific expression noise than regulator genes. These findings show that not just the mean, but also the variance of gene expression level, respond to selection in changing environments, and do so in a network-dependent manner. They further point to gene expression noise as a mechanism for populations to escape extinction by creating phenotypic heterogeneity when faced with environmental changes.

## Introduction

A biological organism is formed through an interaction between the environment and its genome, which together drive how genetic information is expressed into the phenotype. An important component of the gene expression profile, in addition to the mean gene expression level, is the variability of gene expression level between cells and tissues. Although gene expression has been extensively measured using bulk transcriptomics, averaging expression levels over many cells, the advent of single-cell biology allows quantifying the intrinsic variability of expression levels between isogenic cells (Kærn et al., 2005). The cell-to-cell expression variance present in isogenic cells grown in a homogeneous environment is termed gene expression noise and stems from the probabilistic molecular reactions involved in the many steps of the gene expression process. Expression noise is distinct from gene expression plasticity, the variance of the gene expression level induced by different environmental conditions (Sommer, 2020), because the expression variance caused by expression noise is not caused by environmental signaling, but the unavoidable physical nature of diffusion and binding mechanics within the cell. As such, expression noise can only be decreased or increased through different molecular or kinetic mechanisms, for example, by changing gene transcription and translation rate (Hausser et al., 2019), but cannot be fully removed.

Expression noise has been shown to be an evolvable trait partially independent of the expression mean (Duveau et al., 2018; Barroso et al., 2018; Wolf et al., 2015). For example, stabilizing selection at the gene expression level reduces expression noise (Lehner, 2008). Expression noise has been suggested to be beneficial during the adaptation of the mean expression level to a new expression level optimum in genes under directional selection (Duveau et al., 2018). Increasing expression noise as a bet-hedging strategy for adapting to fluctuating environments was experimentally demonstrated in previous studies. Bet-hedging is the phenomenon of increasing phenotypic variation, which decreases the short-term average fitness of the population but increases the long-term fitness in changing environments, allowing it to survive environmental fluctuations (Grimbergen et al., 2015). In the same way, expression noise can create phenotypic heterogeneity in a clonal population and improve its capacity to survive in changing environments. For example, the increased cell-to-cell variability of a signal transduction system in *Escherichia coli* allows growth in case of rapid fluctuations in oxygen availability (Carey et al., 2018). Expression noise of the transcription factor *comK* was shown to drive cell fate determination in *Bacillus subtilis* and enables competency in a proportion of the cell population (Maamar et al., 2007), and increasing the noise increases the response range of the competency circuit (Mugler et al., 2016). A recent study has shown that the increase in gene expression noise at low growth rates confers a fitness advantage to microbes in unpredictable environments and allows them to explore new phenotypes to adapt to unfavourable conditions (de Groot et al., 2023). In addition, several observed properties of gene regulatory networks have been attributed to fluctuating selection in gene networks (Tsuda and Kawata, 2010).

The response of expression noise to different selection scenarios has been extensively studied in individual genes but not in the context of gene networks. Since expression noise has been demonstrated to propagate in gene networks (Pedraza, 2005), it is important to take into account the network background of a gene under selection when investigating the evolution of expression noise. Changing the expression noise of a gene changes the expression noise of downstream elements because noise propagates between genes in a network. As such, the effective expression noise of a gene is the sum of the expression noise caused by gene-specific sequence features (*e.g.* promoter shape (Sigalova et al., 2020)), and the noise propagated from the transcription factors that affect its expression. Studies in several organisms suggest that the expression levels of most genes evolve under stabilizing selection (Gilad et al., 2006; Lehner, 2008), and consequently that low expression noise is maintained. Increasing expression noise in some parts of the network as a response to fluctuating or directional selection would increase noise in neighbouring parts of the network, which might conflict with stabilizing selection acting on other genes and decrease the efficiency of selection. Therefore, adaptation to directional or fluctuating selection at the gene expression level of individual genes might be constrained by their position in the gene network. How expression noise responds to selection in changing environments when the gene in embedded in a regulatory network remains poorly understood.

In this study, we used a computational gene regulatory network model to simulate the evolution of populations of model gene regulatory networks in two different selection scenarios, directional and fluctuating selection acting on gene expression levels, to investigate the adaptability constraints imposed on individual genes by their gene network background. We found that higher expression noise is transiently beneficial under directional selection, while the mean expression level is evolving towards the new expression optimum. Higher expression noise is also beneficial under fluctuating selection. In addition, the adaptation rate and adaptation probability of gene expression mean and noise is not independent of the network background. Namely, regulator genes are less likely to adapt to directional selection than non-regulator genes because the change to their mean expression level impacts the mean expression level of downstream genes. Regulators are also less likely to adapt to fluctuating selection by increasing their expression noise as part of a bet-hedging strategy.

## 1 Methods

To study the evolution of gene expression noise and mean expression level in gene regulatory networks in changing environments, we simulated the evolution of populations of model gene regulatory networks under directional and fluctuating selection.

### 1.1 Gene regulatory network model with evolvable gene expression mean and noise level

We model a gene network of transcription factors (regulator genes) and their targets (effectors, regulated genes). Gene expression is considered as a single stochastic process, *i.e.* we do not distinguish dynamic chromatin accessibility, transcription, posttranscriptional modifications, translation or posttranslational modifications. Instead, we abstract all steps of gene expression into a single process which yields an expression level distribution with a certain mean and variance, enabling us to study the evolution of expression mean and variance in the simplest terms. The expression level in this model can be understood as the protein level, as transcription factors and most target genes need to be expressed as proteins to fulfill their functions.

We used the formalism introduced by Wagner (Wagner, 1996) as a basis to model networks of *n* genes using a regulatory network matrix *W* = (w_ij_)_1≤i≤n, 1≤j≤n_, in which entries *w_ij_* determine the presence, strength, and sign of the effect of gene *j* on the expression level of gene *i*. The sign of the elements of the regulatory network matrix *w_ij_* determines whether the interaction represents an activation or repression of the downstream expression level, and the value determines the strength of the interaction (Figure 1A). The equilibrium expression level of each gene, if it can converge to a steady state, is considered to be the phenotype of the network. A fitness value is then calculated as the distance to a given optimal expression level with a fitness function which penalizes the distance of each gene from the optimal expression level. The model was extended to account for the intrinsic noise of each gene via gene-specific parameters 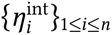 (Puzović et al., 2023) and the basal expression level of each gene 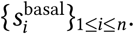

**Figure 1:**
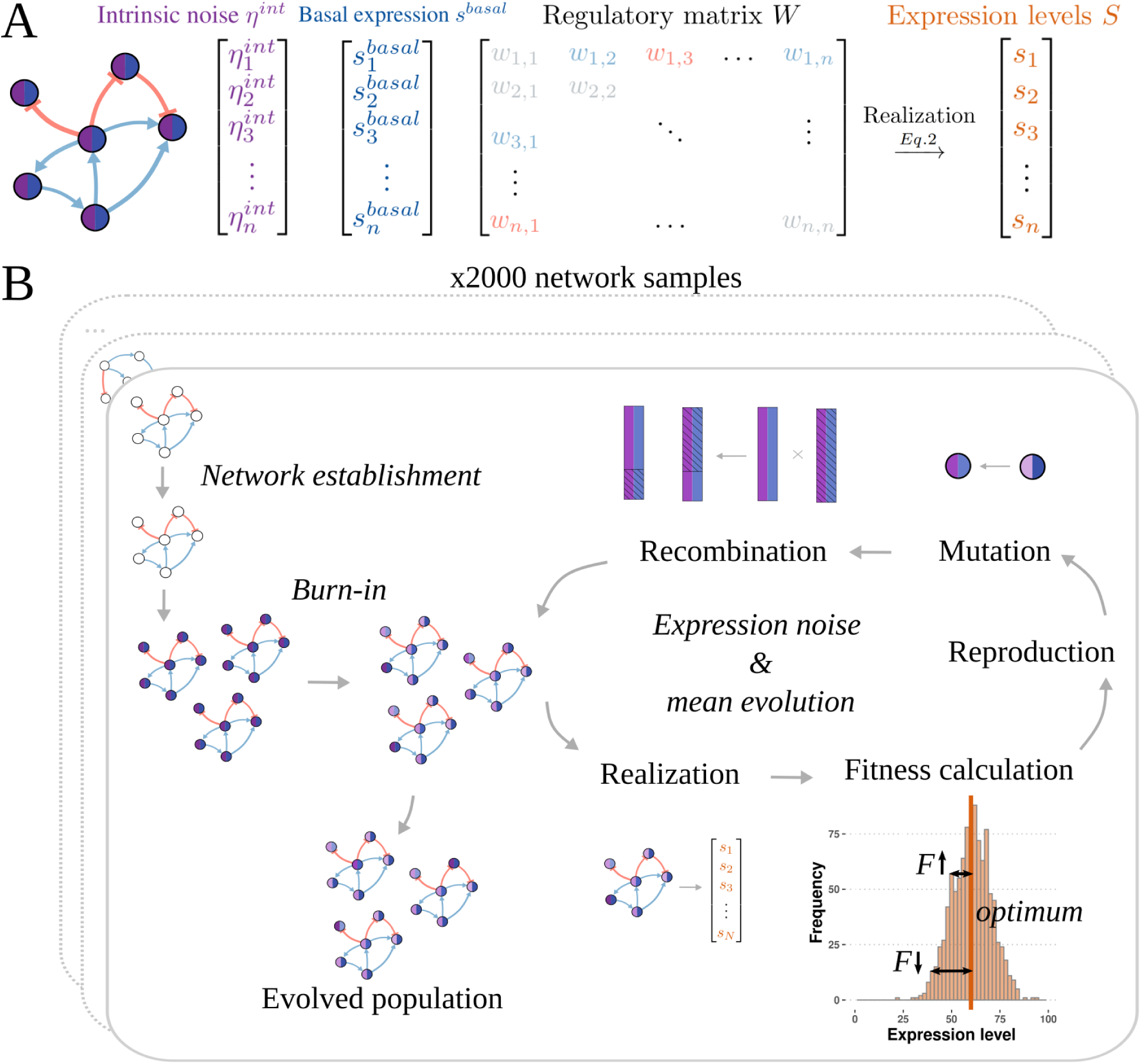
The evolution of gene-specific expression noise was simulated using populations of model gene regulatory networks with mutable levels of gene-specific expression noise under selective and non-selective conditions. A - Gene regulatory network model. The genotype consists of the intrinsic noise vector *η^int^*, basal expression level vector *s*^basal^ and regulatory matrix *W*. The phenotype is given by the state vector *S*, which represents the expression level of each gene in the network. B - Steps of the evolutionary simulation process. Each established network configuration was used as a founding network for the network populations used in the noise evolution simulation. In every generation, genotypes are realized and their fitness assessed. Genotypes are reproduced based on their relative fitness and mutations and recombination events are introduced. The process is repeated for 10,000 generations.

The genotype is realized into the phenotype through a set of difference equations, updating the expression level *s_i_*(*t*) of each gene in every time step *t* using the following rule:

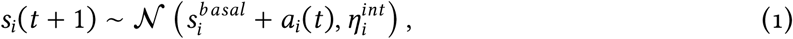

where 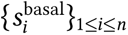 represents the basal expression level of each gene, that is, the constitutive expression level of each gene that is present regardless of the input from the regulatory genes. In this stochastic version, the expression level of each gene in each time step is drawn from a normal distribution, with a mean equal to the sum of the activation rate in the previous time step *a_i_*(*t*) and the basal expression level, and a standard deviation equal to the intrinsic noise parameter. The activation rate is defined as the sum of the effects of all regulators:

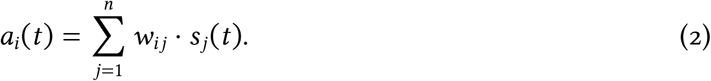

If the value of the expression level drawn is below 0 or above *s_max_* = 100, its value is set to 0 or *s_max_*, respectively, thus constraining the expression level range to an interval (0, 100). The maximum expression level was imposed to prevent the gene expression level from converging to infinity and can be understood as a parameter that reflects both the protein degradation rate and the metabolic limit that prevents the expression level, *i.e.* protein abundance, from increasing above a certain threshold. The intrinsic noise parameter of each gene directly affects the variance of the expression level distribution, however, the expression variance is also directly affected by the variance of upstream transcription factors. The expression variance of each gene is, therefore, a sum of the gene-specific expression noise, which we term intrinsic noise, and the noise propagated from upstream regulators. The expression levels of each gene are updated synchronously in every time step *t* for the *T_r_* time steps (*T_r_* = 50 in this study). The expression level vector {*s_i_*}_1≤*i*≤*n*_ at the final time step *t*_50_ is taken as the phenotype of the individual network. An individual genotype consists of the regulatory network matrix *W*, the vector of intrinsic noise parameters 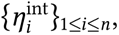 and the vector of basal expression levels 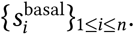

### 1.2 Forward-in-time simulation of expression mean and noise evolution

Populations of model gene regulatory networks were evolved for *T* = 10, 000 generations using an evolutionary algorithm consisting of repeated cycles of phenotype realization, reproduction, mutation and recombination (Figure 1B). First, all individuals in the population had their genotype realized into a phenotype. Next, the fitness of each individual was calculated as a function of the weighted distance of gene expression levels from an optimal gene expression level vector, where weights *ρ*_1≤*i*≤*n*_ are the fitness contribution of each gene:

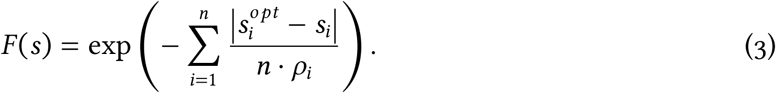

The fitness contribution *ρ* defines the magnitude of the fitness cost resulting from the deviation from the optimum expression level. In this study, we set ∀*i*, *ρ_i_* = 1, defining an equally strong selective pressure on all genes. Individuals in the next generation were randomly sampled from the population in the current generation, with genotype probabilities proportional to their respective fitness values. A constant population size was maintained. Mutations were introduced in the intrinsic noise vector and the basal expression level vectors with a probability of *μ_η_* = 0.005 and *μ_sb_* = 0.005 per gene, respectively. Mutation values for intrinsic noise mutations were drawn from a uniform distribution *U* (0, 200), and mutation values for basal expression level mutations were drawn from a uniform distribution *U* (0, 100). Recombination was implemented by drawing a recombining individual with a probability *r* = 0.05, randomly drawing two genome break points, and exchanging the recombining fragments with another randomly drawn individual in the population. Recombination was carried out independently in noise vectors and basal expression level vectors. The network topology was immutable in the noise evolution simulations: network interactions could neither be lost nor gained, and the sign and strength of the interactions was fixed.

### 1.3 Simulation pipeline

We generated 2,000 unique 40-gene random (Erdős–Rényi) network topology samples with a network density of *d* = 0.05 using the *igraph* package (Csardi and Nepusz, 2006) in R (version 3.6.3, Team (2021)). Each unique network topology sample was used to simulate the evolution of one population of networks, which share the same network configuration but may differ in their basal expression level or expression noise.

Gene regulatory networks generated by randomly assigning the strength of the interactions usually had a similar network-wide gene expression profile; many genes were expressed at the maximum value or were not expressed at all. For a more realistic expression level profile with intermediate gene expression levels, a network establishment step was performed through evolutionary simulations similar to the main noise evolution simulations. In this first step, the strength of the interactions was allowed to mutate, but the overall network topology was fixed; that is, non-null regulatory strengths cannot become zero, and regulatory strengths equal to zero cannot change value. The intrinsic noise of all genes was set to zero and the values for the basal expression levels were drawn from a uniform distribution U(0, *s_max_*) to form the starting population. Both intrinsic noise and basal expression levels were immutable. The target (optimal) expression level for all genes in this step was 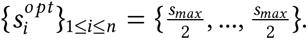 For each of the 2,000 network topologies, a population of *N* = 1, 000 individuals was evolved for *T_est_* = 5, 000 generations. The strength of the regulatory interactions values were drawn from a uniform distribution U(−3, 3) with a mutation probability of *μ_reg_* = 0.001 per link. Networks that had oscillatory gene expression trajectories were removed. The fittest network at the end of the network establishment process had more genes with intermediate expression levels than the network in the starting population, and its equilibrium expression level values were used to set the optimal expression level in the rest of the simulation. The genotype of the fittest network was used as the founding network to create a uniform population for the next step, the burn-in.

Stabilizing selection was applied to all genes in each network population for *T_burn_* = 5, 000 generations until the mutation-selection-drift balance was reached (burn-in step). In this step, the intrinsic noise was set to zero for all genes in the population of *N* = 1, 000 individuals at the beginning of the simulation. The intrinsic noise and basal expression levels were allowed to mutate. Intrinsic noise mutations were drawn from a uniform distribution *U* (0, 200) with a mutation rate of *μ_η_* = 0.01 per gene, and basal expression level mutations were drawn from a uniform distribution *U* (0, 100) with a mutation rate of *μ_η_* = 0.01 per gene. The burn-in phase ensured that all 40 genes in each network had already been evolving under stabilizing selection for long enough that the mutation-selection-drift balance had been reached, and any selection response observed after an environmental shift would be due to directional or fluctuating instead of stabilizing selection. The heterogeneous population of genotypes at the end of the burn-in phase was the starting point for the main simulation, when an environmental shift was applied.

One gene in each network was randomly chosen so that its optimum expression level *s^opt^* was increased or decreased by 20% of *s_max_* relative to its previous value. The remaining 39 genes remained under stabilizing selection, with their respective optimum expression level values unchanged. Two selective scenarios were simulated: directional selection, in which the optimum expression level was either increased or decreased by 20% of *s_max_*, and fluctuating selection, in which the optimum expression level oscillated between an increase and a decrease of 20% of *s_max_* every other generation. The intrinsic noise and basal expression levels were mutable and the network topology was immutable. The populations evolved for *T_evol_* = 10, 000 generations. The evolutionary simulation was replicated 10 times for each selective scenario and the resulting data (changes in phenotypes and genotypes) were averaged over the replicates for each gene in each scenario.

To understand the network background’s specific effect on the evolvability of gene expression in changing environments, we also simulated the evolution of single, isolated genes under directional and fluctuating selection. A dataset of 1,000 genes with a random value of the basal expression level drawn from *U* (20, 80) was used to generate populations of *N* = 1, 000 individuals. Their evolution was simulated using the pipeline described above: each population evolved for *T_burn_* = 5, 000 generations under stabilizing selection, after which directional or fluctuating selection was applied for an additional *T_evol_* = 30, 000 generations. As a control, we also let the populations evolve under stabilizing selection. The network modeling framework presented here can be used to investigate the evolution of expression mean, expression noise, or both at the same time, by setting the mutability of the evolutionary parameters, which will be explored in the Results section. The simulation results were analysed in R v3.6.3 (Team (2021)), using the Mistral Medium 3.5 and Large 4 LLM model as a coding assistant. The gene regulatory network model and evolutionary simulator were implemented in C++ and are freely available at https://github.com/NatashaPuzovic/netlings as release version netlings v2.1.1.

## Results

To study the evolution of gene expression noise and mean expression level in regulatory networks when the optimal expression level under selection is not constant, we simulated model networks evolving under scenarios where the gene expression level is under directional or fluctuating selection. In each network, one focal gene was randomly chosen to have its optimal expression level shifted, while the other genes in the network remained under stabilizing selection. The populations of model networks were evolved with mutable gene intrinsic noise and basal expression level, and a fixed network topology.

To better understand the effect of the network structure, we first analyse the evolution of single, isolated genes and then add gene interactions. In order to distinguish the response of the mean expression level, intrinsic noise, and their co-evolutionary dynamics, we simulated three scenarios: (i) with mutable intrinsic noise, but fixed basal expression level, (ii) with mutable intrinsic noise and basal expression level, and (iii) with mutable basal expression level, but fixed intrinsic noise. We then studied the selection response of the phenotype (mean expression level and variance of expression level) and genotype (intrinsic noise and basal expression level parameters of the genotypes) of the populations evolved under directional or fluctuating selection on the gene expression level.

### 1.4 Gene expression noise transiently increases as a response to directional selection

To understand how gene expression level distributions respond to an environmental shift that imposes directional selection on gene expression level, we simulated the evolution of 1,000 isolated genes. The evolutionary trajectory of one example gene under directional selection in the first scenario (mutable intrinsic noise, but fixed basal expression level) is shown in Fig. 2A-D. The mean expression level did not change as the basal expression level was immutable (Fig. 2A, D). However, the expression variance and intrinsic noise parameters significantly increased (Fig. 2B, C), indicating that noise was selected for and beneficial in genes whose expression level was far from the optimum. In the dataset of 1,000 simulated genes, the average increase in intrinsic noise was significantly higher in genes under directional selection than genes that remained under stabilizing selection (p-value < 2.2 × 10^−16^, Wilcoxon signed-rank test, Fig. 2E).

**Figure 2:**
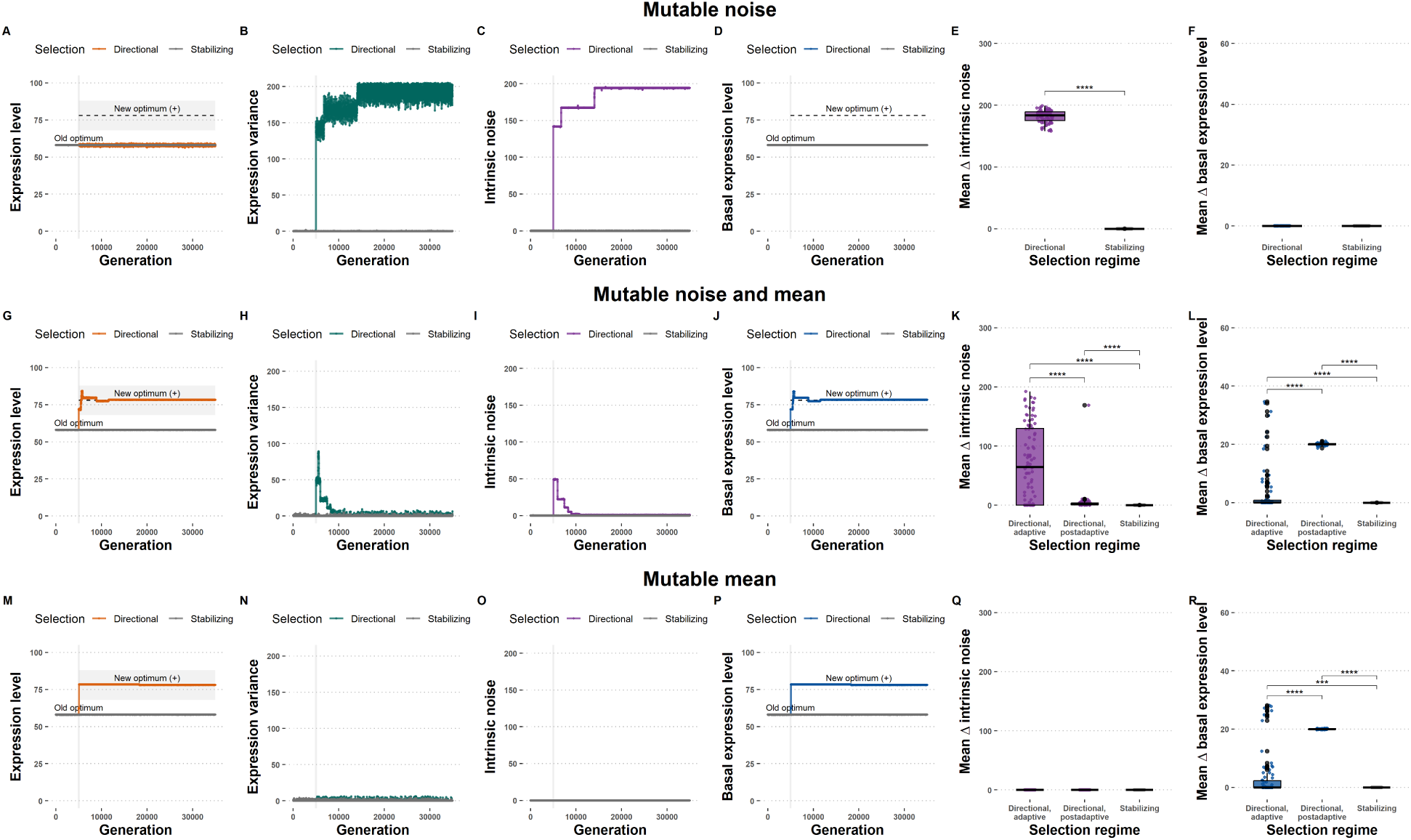
Expression noise is beneficial under directional selection if the mean expression level is fixed, or transiently while the mean is evolving to a new optimum. Three scenarios of evolution under directional selection: mutable noise only (first row **A-E**), mutable noise and basal expression level (second row **F-K**), and mutable basal expression level only (third row, **L-P**). First column **A,F,L**: mean expression level. Second column **B,G,M**: expression level variance. Third column **C,H,N**: intrinsic noise. Fourth column **D,I,O**: basal expression level. Fifth column **E,J**: Average change of intrinsic noise. Sixth column **K, P**: Average change of basal expression levels. Dataset consists of 1,000 genes evolved for 30,000 generations. Asterisks indicate statistical significance of Wilcoxon signed-rank tests: n.s. - p-value > 0.05; * - p-value ≤ 0.05; ** - p-value ≤ 0.01; *** - p-value ≤ 0.001; **** - p-value ≤ 0.0001.

In the second scenario (mutable intrinsic noise and mutable basal expression level), we observed two distinct evolutionary phases. The adaptive phase, during which the basal expression level was evolving towards the new optimum expression level, and the post-adaptive phase, after the basal expression level has reached the optimum expression level and the selection regime switched back to stabilizing selection. The evolutionary trajectory of one example gene is shown in Fig. 2F-I. A co-evolutionary pattern was observed – the expression level variance, determined by the intrinsic noise, was elevated while the basal expression level was evolving towards the new optimum (Fig. 2F, G). Expression noise and intrinsic noise were reduced once the mean expression level reached the optimum expression level (Fig. 2G, H). The average increase of the intrinsic noise during the adaptive phase was significantly higher in genes under directional selection than genes which remained under stabilizing selection (p-value = 1.25 × 10^−10^, Wilcoxon signed-rank test) or genes under directional selection in the post-adaptive phase (p-value = 8.19 × 10^−07^, Wilcoxon signed-rank test) (Fig. 2J). As expected for genes under directional selection, the average change of the basal expression level was higher in genes under directional selection than genes which remained under stabilizing selection in both adaptive (p-value < 2.2 × 10^−16^, Wilcoxon signed-rank test) and post-adaptive phase (p-value < 2.2 × 10^−16^, Wilcoxon signed-rank test) (Fig. 2K).

In the third scenario (mutable basal expression level, but fixed intrinsic noise) we again observed the adaptive and post-adaptive phases of expression mean evolution. The evolutionary trajectory of an example gene is shown in Fig. 2L-O. There is no increase in the expression level variance during the adaptive phase (Fig. 2M), contrary to the increase in expression level variance due to increased intrinsic noise in the second scenario (Fig. 2G, H). Importantly, the lack of a significant increase in expression level variance during the adaptive phase when intrinsic noise cannot evolve indicates that the basal expression level variants segregating in the population did not cause the elevated expression variance signal in the scenario in which both mean and noise could evolve (Fig. 2G). Therefore, the elevated expression variance signal stems from the selectively favoured increase of intrinsic noise during the period in which the mean expression level is evolving towards a new expression optimum.

These results showed that gene expression noise has a fitness benefit if the mean expression level is not near the optimal level, *i.e.* the gene is under directional selection. We simulated the evolution of genes under directional selection, by either increasing or decreasing the optimal expression level. We observed consistent results in the two cases and for concision only reported the results from directional selection with an increased optimal expression level in the main text, while the results from directional selection with a decreased optimal expression level can be found in Section 1 of the Supplementary Information. We observed that expression noise remains elevated under directional selection if the mean expression level is constrained and cannot change. However, if the mean expression level can evolve towards a new optimum, a switch in the selection regime occurs as the population gets closer to the new optimum: the expression level is under stabilizing selection again, and expression noise becomes deleterious. Therefore, expression noise is transiently beneficial while the expression mean is evolving towards a new optimum, but becomes deleterious once it has reached the new peak. If the expression mean cannot evolve, expression noise has a constant fitness benefit. This may happen if the evolution of the mean expression level is constrained by the position of the gene in the regulatory network, which is what we explore in the next section.

### 1.5 Regulator genes in gene regulatory networks are less adaptable to directional selection than target genes

Next, we investigate whether the network background has an effect on the evolvability of genes connected in a regulatory network. The evolutionary trajectory of a gene under directional selection from an example 40-gene network is shown in (Fig 3A-D). Before a directional selection regime was applied, the population was at mutation-selection-drift equilibrium, and the expression level of the focal gene was under stabilizing selection. Consequently, the population-wide mean expression level showed little variation (Fig 3A, MSD balance phase). After the optimal expression level changed, the population mean expression increased toward the new optimum. We categorised genes as adapted if their mean expression level reached optimal expression level *s^opt^* ± 10%*s_max_*. As in the single gene case, we distinguish two evolutionary phases in genes whose expression level responded to directional selection: i) adaptation, during which the expression mean is evolving towards the new optimum, and ii) post-adaptation, after the expression mean has converged toward a new optimum. During these two phases, we track changes in expression variance (Fig 3B), intrinsic noise genotypes (Fig 3C) and basal expression level genotypes (Fig 3D). Expression variance is a reflection of the expression variance caused both by gene-specific (intrinsic) noise and the noise propagated from upstream regulators, if there are any. In the depicted example gene, the gene expression level converged toward the new optimum. During the adaptation phase, the average population-wide expression variance and intrinsic noise parameters increased. Conversely, in the post-adaptation phase, the phenotypic and intrinsic noise decreased to their average values before the optimum shift (Fig 3B, C).

**Figure 3:**
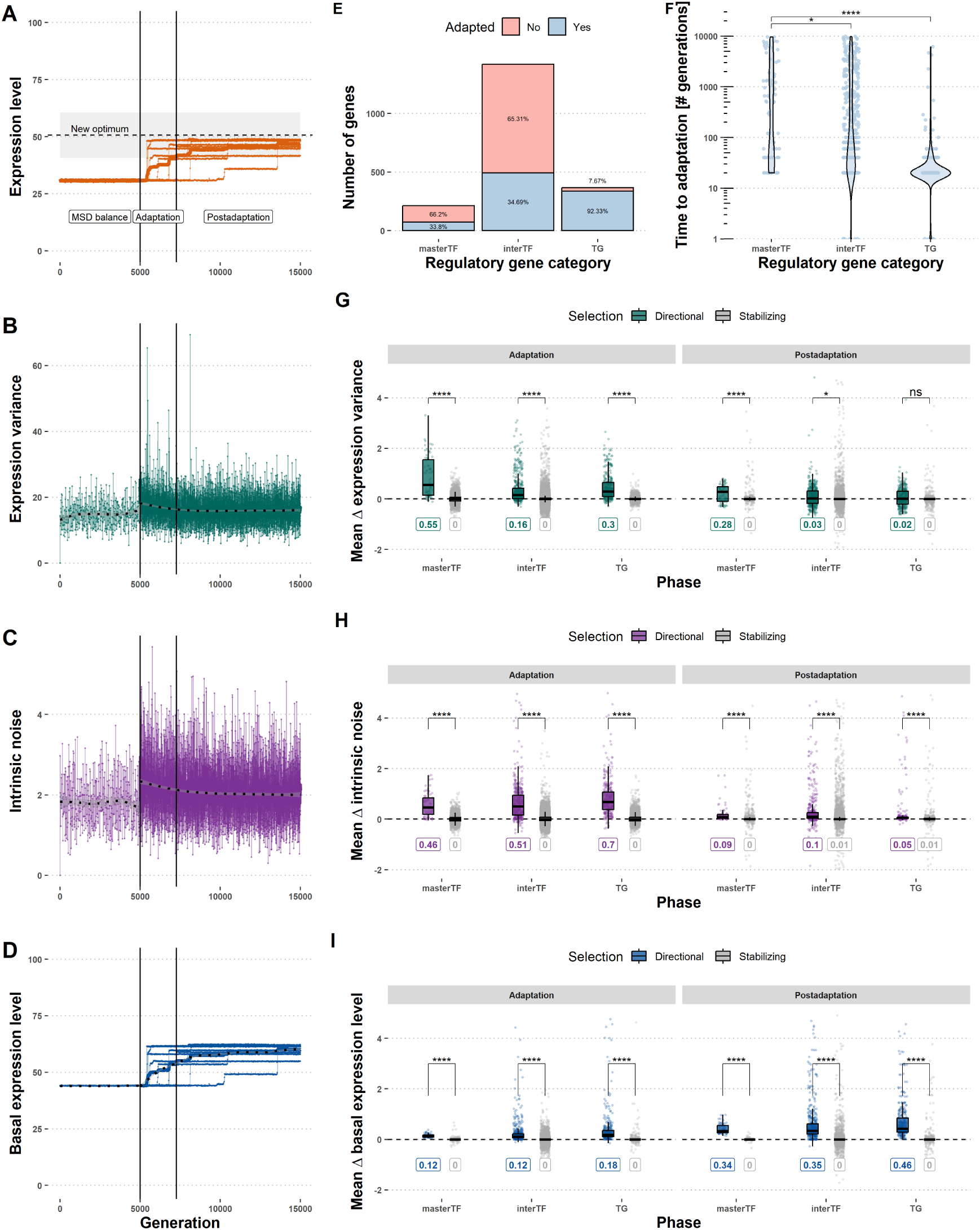
Transcription factors adapt to directional selection on gene expression level less frequently and more slowly than target genes. **A-D** Evolutionary trajectory of an example gene evolved under directional selection (A: mean expression level, B: expression level variance, C: intrinsic noise, D: basal expression level). Black vertical lines indicate the onset of the environmental shift, which imposes the new optimal expression level, and the ending of the adaptive phase, respectively. Dashed horizontal lines indicate the new optimal expression level. **E** Proportion of genes in each regulatory category that responded to directional selection. **F** Time to adaptation in each regulatory category. **G-I** Relative changes of parameters in each regulatory category (G: expression variance, H: intrinsic noise, I: basal expression level). The dataset consists of 2,000 40-gene networks evolved in 10 replicates each for 10,000 generations. Acronyms: MSD balance - Mutation-selection-drift balance. Asterisks indicate statistical significance of Bonferroni-corrected Wilcoxon signed-rank tests. Significance code as in Fig. 2.

Out of 2000 genes under directional selection, 899 (44%) adapted after 10,000 generations. The regulatory function of the selected gene had a significant effect on the probability of adaptation and the time to reach the new optimum. We categorized genes into three regulatory categories based on their regulatory interactions: i) master regulators or transcription factors (masterTFs), which regulate one or more downstream genes but are not regulated by any upstream gene; ii) intermediate regulators or transcription factors (interTFs), which are both regulated and regulate one or more genes; and iii) target genes (TGs), which are regulated by one or more upstream genes, but do not regulate any downstream gene themselves. A higher proportion of target genes adapted to directional selection than intermediate transcription factors and master transcription factors (p-value < 2.2 × 10^−16^, proportion test). Of all genes put under directional selection, 92% of target genes (TG) adapted to the new expression level optimum, contrary to 35% of intermediate transcription factors (interTF) and 34% of master transcription factors (masterTF), indicating a constraint on the evolvability of transcription factor expression levels (Fig 3E).

The time to adaptation, defined as the number of generations until the new optimum expression level is reached, also differed between the three regulatory categories. Adaptation was significantly slower in master transcription factors (p-value < 2.2 × 10^−16^, Wilcoxon signed-rank test) and intermediate transcription factors (p-value < 2.2×10^−16^, Wilcoxon signed-rank test) than in target genes (Fig 3F).

To assess the effect of the interaction strength on the adaptation propensity, we fitted linear models with two node-level network metrics (absolute node instrength and absolute node outstrength), the results of which are summarised in Table 1. Absolute instrength and absolute outstrength had a significant effect on the probability of adaptation under directional selection. Absolute instrength, a metric of the degree to which other genes regulate a gene, had a significant positive effect on the probability of adaptation (Table 1). In contrast, absolute outstrength, a metric of how strongly a gene regulates other genes, had a significant negative effect on the probability of adaptation. There is no significant correlation between the two network metrics (Spearman’s *ρ* = −0.02, p-value = 0.28), but there is a relationship in which genes with high values of instrength have an outstrength of zero, and vice versa, *i.e.* the distributions of both network metrics are zero-inflated (Supplementary Information, Section 2). Therefore, we fitted the same linear models in subsets of each gene regulatory category. In target genes, which have a non-zero value of absolute instrength, and an absolute outstrength of zero, the absolute instrength did not have a significant effect on the probability of adapting to selection (Table 1). In intermediate transcription factors, which have non-zero values for both absolute instrength and outstrength, absolute instrength had a significant positive effect on the probability of adaptation, while absolute outstrength had a significant negative effect on the probability of adaptation, echoing the results from the joint dataset. In master transcription factors, genes which have non-zero values for outstrength, but a zero value for instrength, the absolute outstrength had a significant negative effect on the probability of adaptation. Notably, the negative effect size of the absolute outstrength on the adaptation probability was consistently higher than the positive effect size of the absolute instrength.

**Table 1:** Network centrality metrics affect the adaptation probability and adaptation speed in responding to directional selection on gene expression level.

| Response | Reg. category | Explanatory variable | Beta | p-value<sup>1</sup> |  |
|---|---|---|---|---|---|
| Adaptation probability | All | Absolute instrength | 0.094 | 2.01 × 10<sup>−6</sup> | *** |
|  |  | Absolute outstrength | -0.597 | < 2.2 × 10<sup>−16</sup> | **** |
|  | Target gene | Absolute instrength | -0.002 | 0.975 | n.s. |
|  | Intermediate TF | Absolute instrength | 0.087 | 9.96 × 10<sup>−5</sup> | **** |
|  |  | Absolute outstrength | -0.402 | < 2.2 × 10<sup>−16</sup> | **** |
|  | Master TF | Absolute outstrength | -0.558 | 2.24 × 10<sup>−07</sup> | **** |
| Time to adaptation | All | Absolute instrength | -0.029 | 0.006 | ** |
|  |  | Absolute outstrength | 0.105 | 1.33 × 10<sup>−10</sup> | **** |
|  | Target gene | Absolute instrength | 0.012 | 0.157 | n.s. |
|  | Intermediate TF | Absolute instrength | -0.009 | 0.574 | n.s. |
|  |  | Absolute outstrength | -0.027 | 0.210 | n.s. |
|  | Master TF | Absolute outstrength | 0.212 | 0.008 | ** |

<sup>1</sup> Coefficients and their significance were computed using linear models (see Methods). The dataset consisted of 2,000 populations with unique 40-gene random, which were independently evolved 10 times under selection. Asterisks indicate statistical significance: n.s. - p-value > 0.05; \* - p-value ≤ 0.05; \*\* - p-value ≤ 0.01; \*\*\* - p-value ≤ 0.001; \*\*\*\* - p-value ≤ 0.0001.

A similar pattern was observed for the time to adaptation. Again, the absolute instrength had a significant negative effect on the number of generations until the new optimum was reached (Table 1), while absolute outstrength had a significant positive effect. Absolute outstrength also had a significant negative effect on adaptation speed among transcription factors. In target genes and intermediate transcription factors, the two network metrics did not have a significant effect on adaptation speed.

Lastly, we investigated the response of the expression variance, intrinsic noise, and basal expression level during the adaptation and post-adaptation phases in the genes that managed to adapt to the new expression level optimum. As in the case of isolated genes, genes in networks under directional selection had significantly increased their expression variance (p-value < 2.2 × 10^−16^, Wilcoxon signed-rank test) and intrinsic noise (p-value < 2.2 × 10^−16^, Wilcoxon signed-rank test) during the adaptation phase, relative to their pre-optimum shift and relative to genes that remained under stabilizing selection. A significant increase in expression variance and intrinsic noise was observed in all three gene regulatory categories (Fig 3G, H). After adaptation, the expression variance and intrinsic noise decreased to similar values than before the optimum shift. Genes that remained under stabilizing selection after the directional selection regime was applied to a random gene in their network also showed a statistically significant, but extremely small, increase in average expression variance and intrinsic noise, but not in their basal expression level. This small average increase in expression variance and intrinsic can be explained by their connection to genes under directional selection, whose response to directional selection would affect local genes through noise propagation and inflate the expression variance of the network. As expected, genes under directional selection increased their basal expression level (Fig 3I), compared to genes under stabilizing selection.

These results show that the position of the gene in the gene regulatory network imposes constraints on the evolvability of its expression level in the face of one-time environmental shifts that would impose directional selection. Namely, transcription factors are less likely to adapt to a new optimal expression level and do so more slowly. Next, we explore what happens in genes that experience periodic environmental shifts, which impose a fluctuating selection regime on the gene expression level.

### 1.6 Gene expression noise is increased as a response to fluctuating selection

To investigate whether elevated gene expression noise could be maintained in constantly changing environments, we first simulated the evolution of isolated genes under fluctuating selection on the gene expression level. This fluctuating selection was implemented by imposing switches of the optimum expression level every other generation so that the population did not have time to adapt to the new optimum before a new one occurred. As in the directional selection case, we first report results from the simulations of single, isolated genes and then study the case where the gene is part of a regulatory network. For isolated gene evolution simulations, we simulated the same three scenarios as in section 1.4: i) with mutable intrinsic noise only, ii) with mutable both intrinsic noise and basal expression level, and iii) with mutable basal expression level only. The evolutionary trajectory of one example gene under fluctuating selection in the first scenario is shown in Fig. 4A-D. The optimal expression level alternates every second generation, and the mean expression level does not adapt to any single optimum (Fig. 4A), as the basal expression level is fixed in this scenario (Fig. 4D). However, the expression variance (Fig. 4B) and intrinsic noise (Fig. 4C) are significantly increased after the application of fluctuating selection. In the entire dataset of 1,000 simulated genes, the average change in intrinsic noise relative to pre-selection values was significantly higher in genes under fluctuating selection than in genes that remained under stabilizing selection (p-value < 2.2 × 10^−16^, Wilcoxon signed-rank test) (Fig. 4E).

**Figure 4:**
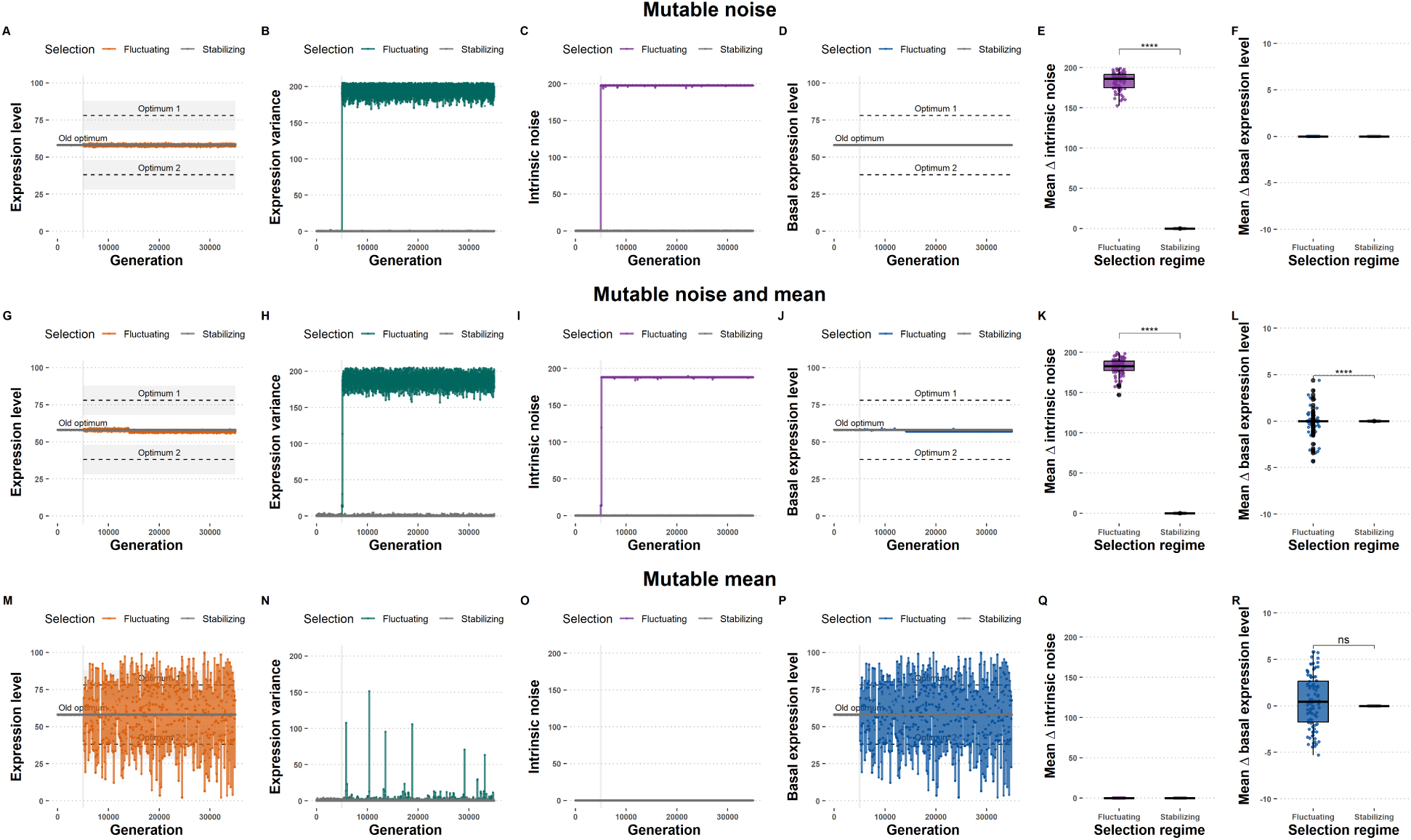
Expression noise is selectively beneficial under fluctuating selection. Three scenarios of evolution under fluctuating selection: mutable noise only (first row **A-E**), mutable noise and basal expression level (second row **F-K**), and mutable basal expression level only (third row, **L-P**). First column **A,F,L**: mean expression level. Second column **B,G,M**: expression level variance. Third column **C,H,N**: intrinsic noise. Fourth column **D,I,O**: basal expression level. Fifth column **E,J**: Average change of intrinsic noise. Sixth column **K,P**: Average change of basal expression levels. The dataset consists of 1,000 genes evolved for 30,000 generations. Asterisks indicate statistical significance of Wilcoxon signed-rank tests. Significance code as in Fig. 2.

Allowing the basal expression level to evolve (second scenario, mutable intrinsic noise and mutable basal expression level) did not alter the observed pattern of increased intrinsic noise after the fluctuating selection regime was applied. The evolutionary trajectory of one example gene is shown in Fig. 4F-I. The mean and basal expression levels did not change significantly under fluctuating selection (Fig. 4F, I), even though the basal expression level was mutable in this scenario.

Since fluctuating selection imposes a new optimal expression level every other generation, it is not beneficial for the population to evolve towards any single optimum. Because the evolution of the mean expression level is constrained, the increased expression noise is maintained as a response to fluctuating selection, as in the case of single genes under directional selection with an immutable mean expression level (Fig. 2E). In the dataset of 1,000 genes, the average change in intrinsic noise relative to pre-selection values was significantly higher in genes under fluctuating selection than in genes that remained under stabilizing selection (p-value < 2.2 × 10^−16^, Wilcoxon signed-rank test) (Fig. 4J). The average change of basal expression level was significantly different between genes under fluctuating and stabilizing selection (Fig. 4K), although the difference is negligibly small.

In the third scenario, in which the basal expression level was mutable but the intrinsic noise was not, the mean expression level and the basal expression level showed large fluctuations between the two expression level optima (Fig. 4L, O). However, the expression variance (Fig. 4M) did not increase as in the second scenario, in which the mean and noise could evolve. This suggests that the increase of expression variance in the second scenario (mutable intrinsic noise and basal expression level) cannot be attributed to the population heterogeneity of basal expression level genotypes while the population was evolving towards a new peak. Instead, the increase in expression variance in the second scenario reflects the increase in intrinsic noise as a fitness benefit in response to fluctuating selection at the gene expression level.

The increase in intrinsic noise after the new selection regime was applied could also be explained by a relaxation of purifying selection. Namely, the drastic environmental shift or alternating environmental shifts throw the population far from its adaptive peak and drastically reduce the average fitness of the population. If the fitness landscape is significantly rugged, the population may be trapped in regions of very low fitness, and selection may be impeded. More deleterious mutations may accumulate, *i.e.* higher intrinsic noise variants would fail to be selected against and would not be removed by selection, and the population-wide intrinsic noise would increase. However, if that were the case, these mutations would be expected to segregate at low frequency, resulting in a population-wide heterogeneity with a mixture of low- and high-noise variants, which is not observed in our study. After the environmental shift occurs, the expression variance and intrinsic noise increase consistently (Fig. 4E, J), and the populations after fluctuating selection are quite homogeneous in high intrinsic noise genotypes. Therefore, the increase in noise is the result of a selective advantage, not a spurious signal from a relaxation of purifying selection against noise.

These results demonstrate that gene expression noise is selectively favoured in genes under fluctuating selection on gene expression level, which can be understood as the evolutionary emergence of a bet-hedging strategy. Lastly, we examine how the network background of the gene under fluctuating selection affects its rate of evolution.

### 1.7 Target genes respond more often and more strongly to fluctuating selection than regulator genes

We investigate whether the background of the gene regulatory network has an effect on the evolvability of genes under a fluctuating selection regime. The evolutionary trajectory of a gene under fluctuating selection from an example 40-gene network is shown in Fig 5A-D. In the mutation-selection-drift balance phase, before fluctuating selection was applied, the expression level of the focal gene was under stabilizing selection. Consequently, the population-wide mean expression level shows little variation (Fig 5A, MSD balance). The mean expression level persisted after fluctuating selection was applied. However, the expression variance (Fig 5B) and the intrinsic noise (Fig 5C) increased in response to fluctuating selection, as in the case of an isolated gene under fluctuating selection.

**Figure 5:**
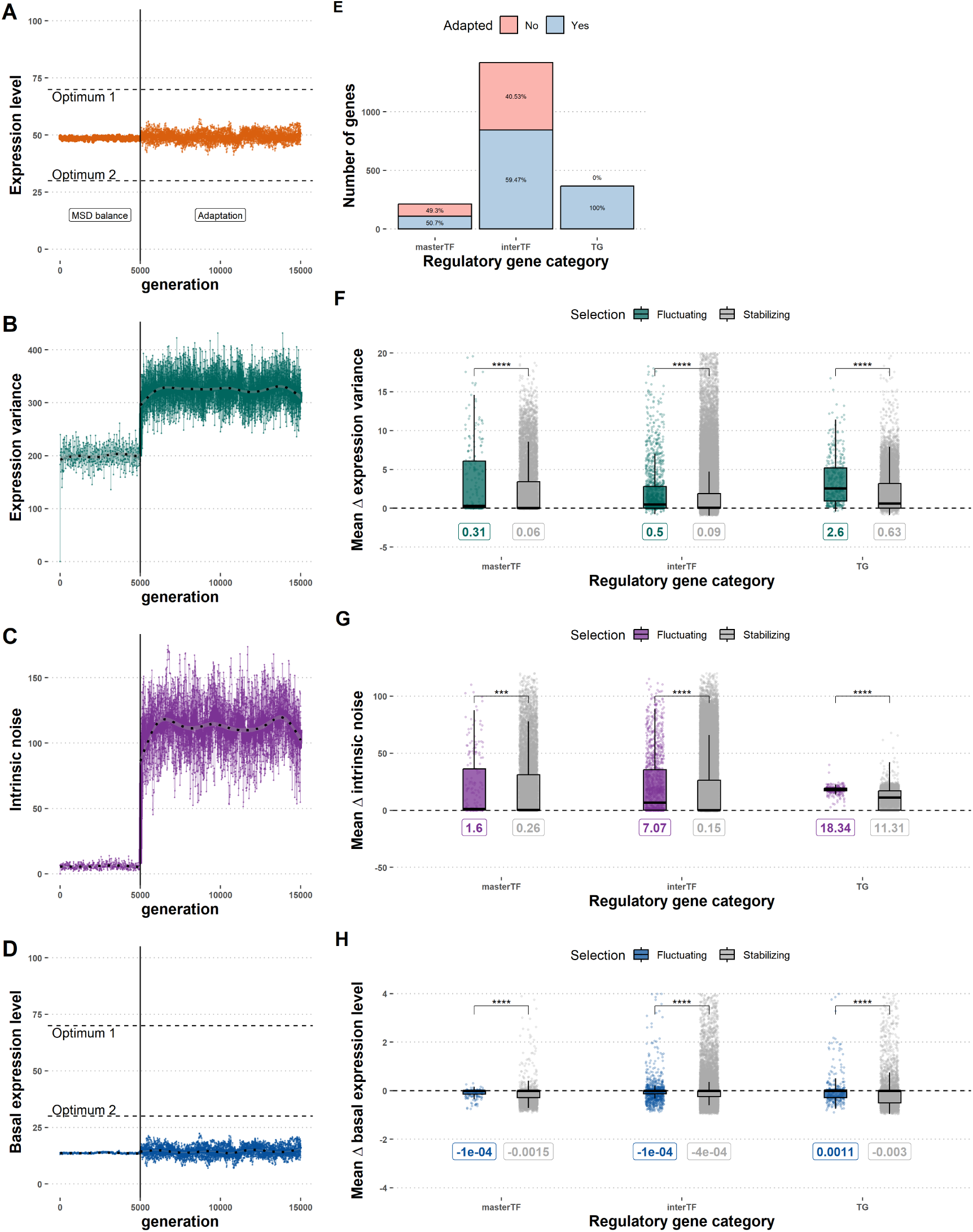
Transcription factors adapt to fluctuating selection on gene expression level less frequently and less strongly than target genes. **A-D** Evolutionary trajectory of an example gene evolved under fluctuating selection (A - mean expression level, B - expression level variance, C - intrinsic noise, D - basal expression level). Black vertical lines indicate the onset of the environmental shift into a fluctuating selection regime. Dashed horizontal lines indicate the optimal expression levels in the two alternating environments. **E** Proportion of genes in each regulatory category that responded to fluctuating selection with an at least twofold increase of their intrinsic noise. **F-H** Relative changes of parameters in each regulatory category (F: expression variance; G: intrinsic noise; H: basal expression level). The dataset consists of 2,000 40-gene networks evolved in 10 replicates for 10,000 generations. Acronyms: MSD balance - Mutation- selection-drift balance. Asterisks indicate statistical significance of Bonferroni-corrected Wilcoxon signed-rank tests. Significance code as in Fig. 2.

The adaptation of gene expression in the case of fluctuating selection is not an adaptation of the mean expression level, but of the expression noise, since the mean expression level is prevented from evolving to any optimum by its frequent switches. Therefore, we classified genes as “adapted” to fluctuating selection if their relative change of intrinsic noise parameters was higher than 1, which represents a twofold increase of intrinsic noise relative to pre-selection switch values. The proportion of genes differed significantly between the three gene regulatory categories (p-value < 2.2 × 10^−16^, proportion test). Of all genes subjected to fluctuating selection, 100% of target genes (TGs) increased their intrinsic noise more than twofold, compared to 59% of intermediate transcription factors (interTF) and 50% of master transcription factors (masterTF) (Fig 5E).

To study the effect of the interaction strength on the noise adaptation probability, we fitted linear models with absolute node instrength and absolute node outstrength as node-level network metrics. The results are summarised in Table 2. In the joint dataset, the absolute instrength had a significant positive effect on the adaptation probability, while absolute outstrength had a significant negative effect on the probability of adaptation. The absolute outstrength also had a significant negative effect on the probability of adaptation in intermediate transcription factors and in master transcription factors (Table 2).

**Table 2:** Network centrality metrics affect the probability of responding to fluctuating selection on gene expression level by increasing intrinsic gene expression noise, and the strength of the response.

| Response | Reg. category | Explanatory variable | Beta | p-value<sup>2</sup> |  |
|---|---|---|---|---|---|
| Noise adaptation probability (signif. increase of noise under fluctuating selection) | All | Absolute instrength | 0.070 | 0.0003 | *** |
|  |  | Absolute outstrength | -0.270 | < 2.2 × 10<sup>−16</sup> | **** |
|  | Target gene | Absolute instrength | NA<sup>1</sup> | *NA*<sup>1</sup> | NA<sup>1</sup> |
|  | Intermediate TF | Absolute instrength | 0.035 | 0.094 | n.s. |
|  |  | Absolute outstrength | -0.141 | 1.1 × 10<sup>−8</sup> | **** |
|  | Master TF | Absolute outstrength | -0.158 | 0.014 | * |

<sup>1</sup> All target genes adapted, so the effect of absolute instrength on the adaptation probability could not be assessed. <sup>2</sup> Coefficients and their significance were computed using linear models (see Methods). The dataset consisted of 2,000 populations with unique 40-gene random, which were independently evolved 10 times under selection. The change of intrinsic noise of each gene was calculated as the average change of the intrinsic noise parameter during the evolutionary simulation and summarized as the mean over all replicates in each scenario. Asterisks indicate statistical significance: n.s. - p-value > 0.05; \* - p-value ≤ 0.05; \*\* - p-value ≤ 0.01; \*\*\* - p-value ≤ 0.001; \*\*\*\* - p-value ≤ 0.0001.

Finally, we looked into the evolutionary trajectory of the expression variance, intrinsic noise and basal expression level of genes in gene regulatory networks undergoing fluctuating selection. In the entire dataset, consisting of 80,000 genes from 2,000 40-gene network topologies evolved in 10 replicates, genes under fluctuating selection had a significantly higher increase in expression variance than genes in the rest of the gene network that remained under stabilizing selection (p-value < 2.2 × 10^−16^, Wilcoxon signed-rank test). Genes subject to fluctuating selection also had a higher increase in intrinsic noise than genes under stabilizing selection (p-value < 2.2 × 10^−16^, Wilcoxon signed-rank test).

The increased expression variance and the intrinsic noise trend were observed in all three gene regulatory categories (Fig 5F-H). However, the degree of increase of the expression variance and intrinsic noise under fluctuating selection was dependent on the position of the gene in the gene regulatory network. Specifically, target genes under fluctuating selection had a significantly higher increase in expression variance than intermediate transcription factors (p-value < 2.2 × 10^−16^, Wilcoxon signed-rank test) and master transcription factors under fluctuating selection (p-value < 2.2 × 10^−16^, Wilcoxon signed-rank test) (Fig 5F). Correspondingly, target genes under fluctuating selection had a higher increase in intrinsic noise than intermediate transcription factors (p-value < 2.2 × 10^−16^, Wilcoxon signed-rank test) and master transcription factors (p-value < 2.2 × 10^−16^, Wilcoxon signed-rank test) (Fig 5G). These results indicate that target genes are less evolutionarily constrained to increase their expression noise as a response to fluctuating selection on gene expression level.

Notably, many genes that remained under stabilizing selection also significantly increased their expression variance and intrinsic noise after another gene in the same network was put under fluctuating selection. In fact, the average change in expression variance and intrinsic noise for genes that remained under stabilizing selection was significantly greater than zero, although still significantly lower than for genes under fluctuating selection. This effect can be explained by the propagation of noise within gene regulatory networks; that is, genes propagate their expression noise downstream, and increasing the intrinsic noise of one gene would increase noise propagated to downstream elements in the network. If a gene is a transcription factor and responds to selection by increasing its intrinsic noise, its increased expression variance would propagate to downstream genes, increasing their expression variance, as well as the overall expression variance in the network. Similarly, if a target gene is under fluctuating selection, it can increase its expression variance by increasing its intrinsic noise, or by increasing the intrinsic noise of any upstream element, which would then propagate the increased noise to the focal gene. These examples showcase that the evolution of gene expression noise, and gene expression level, is not independent among genes in a gene network. Furthermore, an adaptive property of gene expression, such as expression noise, for a gene under selection can come not from the intrinsic gene properties (*cis* regulatory elements), but from a gene to which it is connected to in the network (*trans* regulatory elements). On the other hand, the same mechanism can prevent a transcription factor from increasing noise to respond to fluctuating selection, as it would propagate noise to other genes in the network, which would be deleterious if other parts of the network are under stabilizing selection. Target genes do not have the same constraint – as they do not have downstream elements – and are free to adapt to fluctuating selection by increasing their intrinsic expression noise.

## Discussion

We investigate how natural selection in changing environments affects the levels of gene expression mean and noise in gene regulatory networks. We hypothesize that (i) high levels of expression noise may be favoured after an environmental shift because of the increased proportion of fit phenotypes in the population and (ii) genes differ in adaptability because of the differential constraints resulting from the structure of the regulatory network. To test these hypotheses, we developed a gene regulatory network evolution model with evolvable basal expression level and intrinsic expression noise. We simulated the evolution of gene expression mean and noise in populations under directional and fluctuating selection. We found that expression noise was transiently increased during adaptation but became counter-selected as the population reached the new optimum and the selection regime switched back to stabilizing selection. Maintaining higher levels of expression noise requires a regime of fluctuating selection where optima are frequently changing, before the system enters a stabilizing selection phase. Importantly, regulator genes had a lower probability of responding to selection and responded less strongly than non-regulator genes, indicating a constraining effect of the gene regulatory network on the adaptability of its constituent genes. In the following, we discuss the implications of these results.

### Expression noise and adaptive walks

Our model implements two types of mutation, affecting the basal expression level and each gene’s expression noise, respectively. In this framework, the two types of mutations are independent and experimental evidence shows that expression mean and noise can be decoupled (Metzger et al., 2015; Duveau et al., 2018). We showed that both mutation types are favoured when the expression level is away from the optimum: mutations affecting the basal expression level bring the population closer to the fitness peak, while mutations increasing the expression noise may increase the proportion of fit phenotypes generated by chance. However, the putative advantage of the second type of mutation decreases as the genotypes in the population get closer to the optimal expression levels. High noise levels, therefore, can only persist if the mean expression level is kept away from the optimum level, as we showed with our toy model where the basal expression level was prevented from mutating. This situation also arises in our fluctuating selection regime, where optimal expression levels alternate every generation, preventing the mean expression level from adapting. These results explain why genes involved in the immune system typically display high levels of expression noise (Shalek et al., 2013), which may result from the optimal expression level being determined by constantly changing pathogens. More generally, we posit that high levels of expression noise should be generally expected in any gene involved in an evolutionary arms race.

Although higher expression noise levels do not bring populations closer to the fitness peak and may only be transiently advantageous, mutations increasing the noise level may still prove crucial to the adaptation process. High levels of expression noise increase the phenotypic heterogeneity in the population, ensuring that a certain proportion of the population has a fit phenotype, whichever environmental conditions it finds itself under; an evolutionary strategy known as bet-hedging (Grimbergen et al., 2015). Bet-hedging may allow the survival of the population by allowing the production of individuals with adaptive phenotypes by chance, triggering an evolutionary rescue mechanism in which a declining population manages to survive an environmental shift and recover (Bell, 2017). A parallel can be drawn with bacterial hypermutator strains, which evolve higher rates of genetic mutations and emerge in stressful, fluctuating environments (Matic, 2019). Like expression noise, mutation rate is a second-order trait, *i.e.* it does not change the phenotype directly but governs the rate at which phenotypic variation is generated, and can therefore be favoured when environments change faster than the mean phenotype can adapt. The parallel has limits, however: hypermutability typically arises from defects in DNA repair and carries a long-term cost in deleterious load, whereas expression noise is a tunable regulatory parameter that can be selected up or down without permanent genomic consequence.

In this study’s simulation framework, the population size is kept constant; the individuals are reproduced into the next generation by sampling until the fixed population size is reached. Consequently, the population never goes extinct, even if the individuals have extremely low fitness, and the population will have time to accumulate potentially beneficial mutations and adapt. Conversely, natural populations may go extinct because of mutation load due to the population being far from its fitness optimum. The evolution of bet-hedging might be more apparent if the populations that did not evolve higher levels of expression noise went extinct instead of being propagated to a constant population size. The simulation framework used here can be modified to include the possibility of extinction events by adding a variable population size and a viability threshold. Extending the model in this way would shed light on the putative role of expression noise as an evolutionary rescue mechanism.

### Network-driven epistasis

Adaptive processes have been analysed using the framework of adaptive landscapes, first introduced by Sewall Wright (Wright, 1932), where genotype populations “walk” through a landscape consisting of fitness peaks and valleys, corresponding to genotype configurations that are more or less fit to the given environment. The adaptability of a population, that is, its capacity to reach a certain fitness peak and the rate at which it does so, depends on many factors, such as population genetics parameters (e.g. effective population size), the initial frequency of adaptive alleles, and the genetic architecture of the selected trait (Olson-Manning et al., 2012). A known factor that may slow adaptation is the non-additivity of mutational effects, or *epistasis*, which generates so-called rugged landscapes (Bank, 2022). One epistasis component results from the gene network, where the expression levels of (directly or indirectly) connected genes are not independent. Here, we show that network-driven epistasis leads to differential adaptability of connected genes. Genes with a large regulatory output are less likely to get closer to a new optimum value as their expression may pull multiple other genes away from their respective optima. The potential benefit of increasing their expression noise may also not compensate for the cost of the increased propagated noise to target genes under stabilizing selection. Reducing the number and strength of interactions between genes, *i.e.* reducing the connectivity of the gene regulatory network, would reduce epistasis and thereby increase adaptability to potential selective pressure in the future. In fact, the real biological gene regulatory networks are sparse (Leclerc, 2008), and network sparsity was shown to be an emerging property resulting from optimising the explorability of new phenotypes (Busiello et al., 2017).

In a recent study, Burban et al. (2022) used a Wagner gene network evolution model and showed that domestication, here implemented as a strong directional selection combined with a bottleneck, led to a strong rewiring of the gene regulatory network. More specifically, they observed that domestication resulted in an increase in the number of connections. Several factors may explain the apparent discrepancy between our conclusions and the results of Burban et al. (2022). First, the model of Burban et al. (2022) does not implement intrinsic expression noise while the network structure is fixed in the model we introduced. In particular, the Burban et al. (2022) model did not implement any cost to increased complexity, while we showed that noise propagation imposes a strong constraint on gene evolution, in a network centrality-dependent manner. Adding expression noise to the model of Burban et al. (2022) offers an interesting perspective to understand the impact of domestication (and, more generally, adaptation) on the evolution of regulatory interactions. Ultimately, while computationally challenging, an extended model in which basal expression levels, intrinsic expression noise, and gene interaction evolve would provide insights into the evolution of network complexity in relation to adaptation.

## Acknowledgments

The authors would like to thank Maud Tenaillon and Arnaud Le Rouzic for discussion, Arne Traulsen and Tal Dagan for the helpful suggestions throughout the project. NP was funded by the International Max Planck Research School for Evolutionary Biology (IMPRS Evol Biol).

## Author contributions

**NP**: Conceptualisation, Data collection, Formal analysis, Investigation, Methodology, Software, Visualisation, Writing – original draft, Writing - Review & Editing

**JYD**: Conceptualisation, Methodology, Supervision, Writing - Review & Editing

## Ethics declarations

## Data availability

The scripts, R notebooks and other parameters necessary to rerun the simulations conducted in this study and reproduce the statistical analyses and figures are available at https://github.com/ NatashaPuzovic/supplementarydata_adaptivenoise. The simulation results data can be found at https://doi.org/10.5281/zenodo.10228282.

