## Supplementary Information for "Network-level selection constrains the adaptive evolution of gene expression noise in regulatory genes in changing environments"

**Authors and Affiliations:** Nataša Puzović[1,2*✉], Julien Y. Dutheil[1✉]

**1** Molecular Systems Evolution Research Group, Max Planck Institute for Evolutionary Biology, Plön, Schleswig-Holstein, Germany

**2** Computational Regulatory Genomics Research Group, Max Delbrück Center for Molecular Medicine, Berlin, Germany

### 1 Directional selection (decreased optimum)

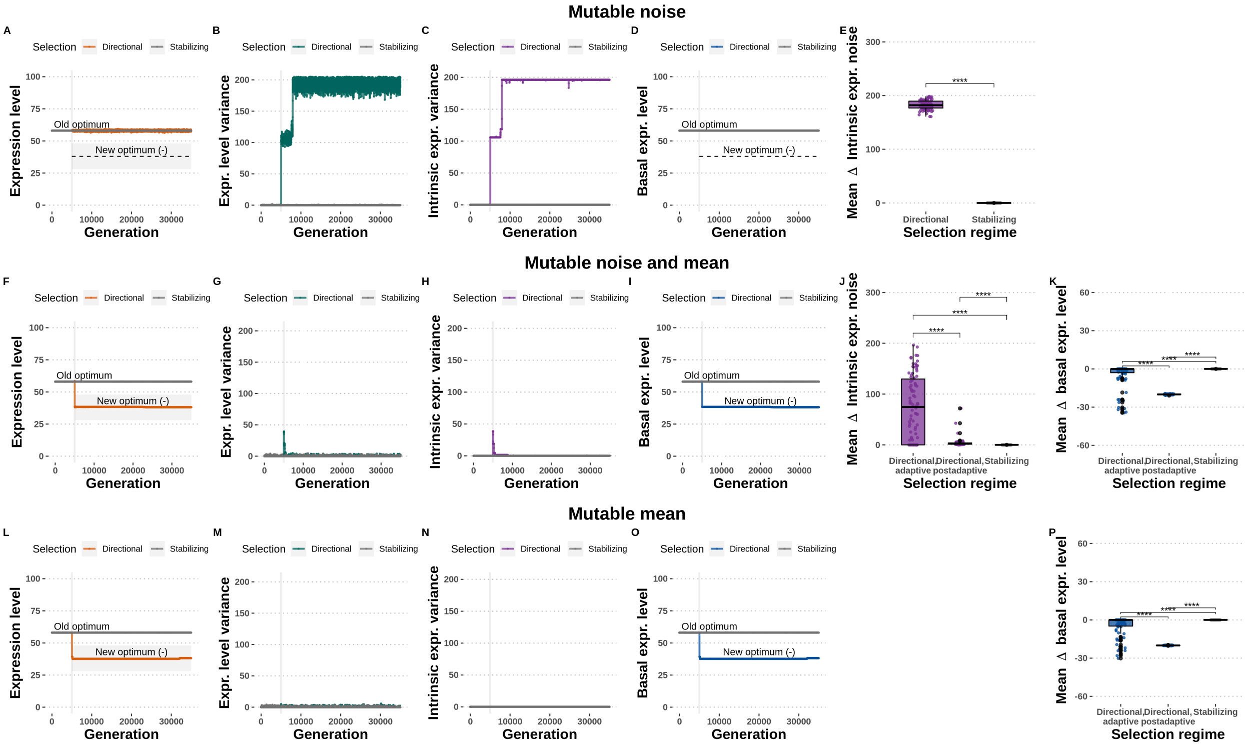


Figure S1: **Expression noise is beneficial under directional selection if the mean expression level is fixed, or transiently while the mean is evolving to a new optimum.** Three scenarios of evolution under directional selection: mutable noise only (first row **A-E**), mutable noise and basal expression level (second row **F-K**), and mutable basal expression level only (third row, **L-P**). First column **A,F,L**: mean expression level. Second column **B,G,M**: expression level variance. Third column **C,H,N**: intrinsic noise. Fourth column **D,I,O**: basal expression level. Fifth column **E,J**: Average change of intrinsic noise. Sixth column **K, P**: Average change of basal expression levels. Dataset consists of 1,000 genes evolved for 30,000 generations. Asterisks indicate statistical significance of Wilcoxon's tests: n.s. - p-value $> 0.05$; * - p-value $\leq 0.05$; ** - p-value $\leq 0.01$; *** - p-value $\leq 0.001$; **** - p-value $\leq 0.0001$.

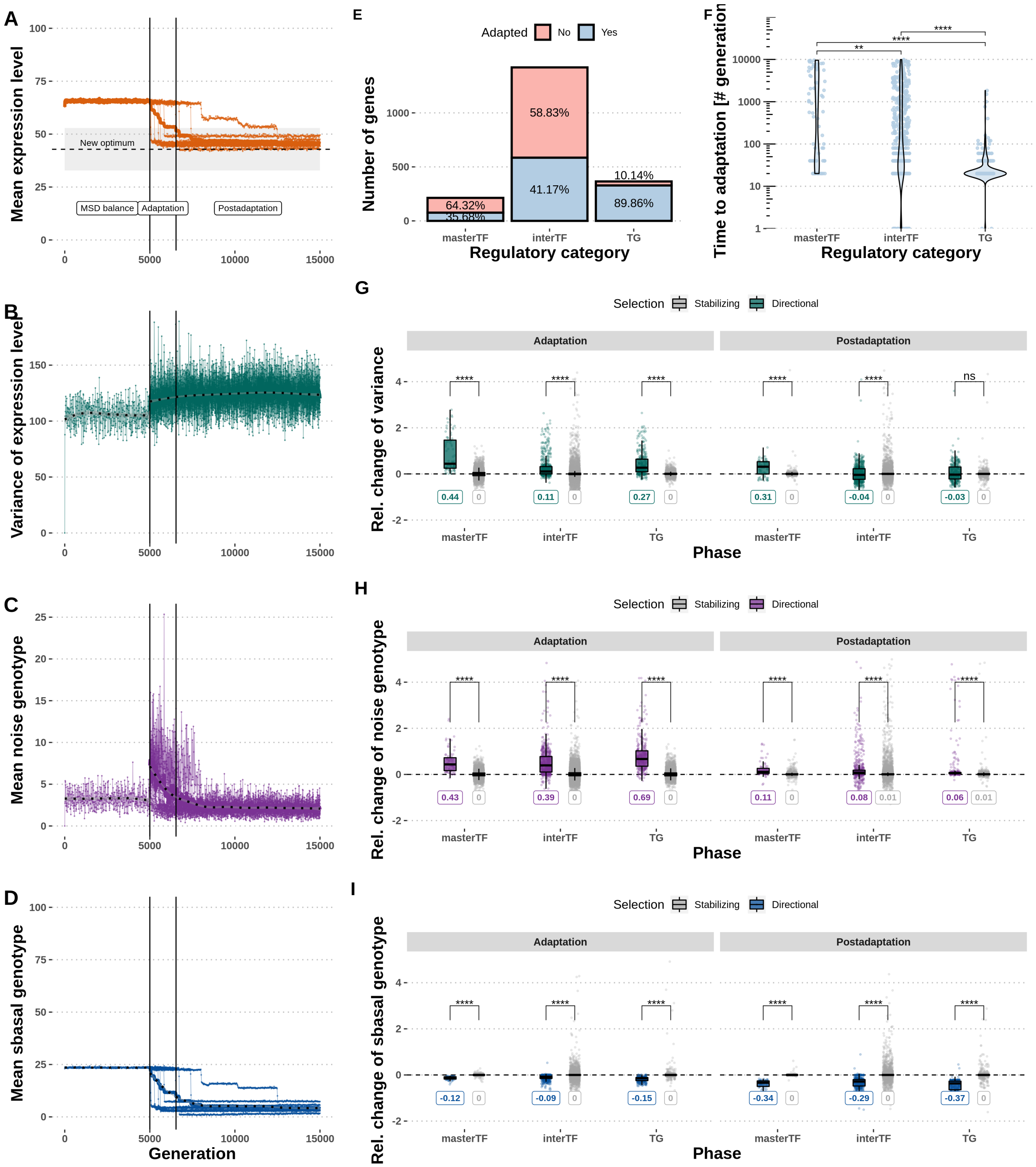


Figure S2: **Adaptation to change in optimal gene expression level in a gene regulatory network.** **A-D** Evolutionary trajectory of an example gene evolved under directional selection (A: mean expression level, B: expression level variance, C: intrinsic noise, D: basal expression level). Black vertical lines indicate the start of the expression level shift, and the ending of the adaptive phase, respectively. Dashed horizontal lines indicate the new optimal expression level. **E** Proportion of genes in each regulatory category that responded to directional selection. **F** Time to adaptation in each regulatory category. **G-I** Relative changes of parameters in each regulatory category (G: phenotypic noise, H: intrinsic noise, I: basal expression level). The dataset consists of 2,000 40-gene networks evolved for 10,000 generations. Acronyms: MSD balance - Mutation-selection-drift balance. Asterisks indicate statistical significance of Wilcoxon's tests. Significance code as in Fig. S1.

Table S1: **Network centrality metrics affect the probability of responding to directional selection on gene expression level by adapting to the new optimal expression level, and the speed of adaptation in generation time.**

| Response | Reg. category | Explanatory variable | Beta | p-value[1] | |
|---|---|---|---|---|---|
| Expression level adaptation probability | All | Absolute instrength | 0.094 | $7.96 \times 10^{-7}$ | **** |
| | | Absolute outstrength | -0.485 | $< 2.2 \times 10^{-16}$ | **** |
| | Target gene | Absolute instrength | 0.011 | 0.879 | n.s. |
| | Intermediate TF | Absolute instrength | 0.078 | 0.0003 | *** |
| | | Absolute outstrength | -0.318 | $< 2.2 \times 10^{-16}$ | **** |
| | Master TF | Absolute outstrength | -0.796 | $8.62 \times 10^{-10}$ | **** |
| Expression level adaptation speed | All | Absolute instrength | -0.067 | $1.18 \times 10^{-10}$ | **** |
| | | Absolute outstrength | 0.088 | $6.65 \times 10^{-9}$ | **** |
| | Target gene | Absolute instrength | 0.009 | 0.155 | n.s. |
| | Intermediate TF | Absolute instrength | -0.065 | $1.16 \times 10^{-5}$ | **** |
| | | Absolute outstrength | -0.012 | 0.542 | n.s. |
| | Master TF | Absolute outstrength | 0.134 | 0.143 | n.s. |

[1] Coefficients and their significance were computed using linear models (see Methods). The dataset consisted of 2,000 populations with unique 40-gene random, which were independently evolved 10 times under selection. Asterisks indicate statistical significance: n.s. - p-value > 0.05; * - p-value ≤ 0.05; ** - p-value ≤ 0.01; *** - p-value ≤ 0.001; **** - p-value ≤ 0.0001.

### 2 Network metrics

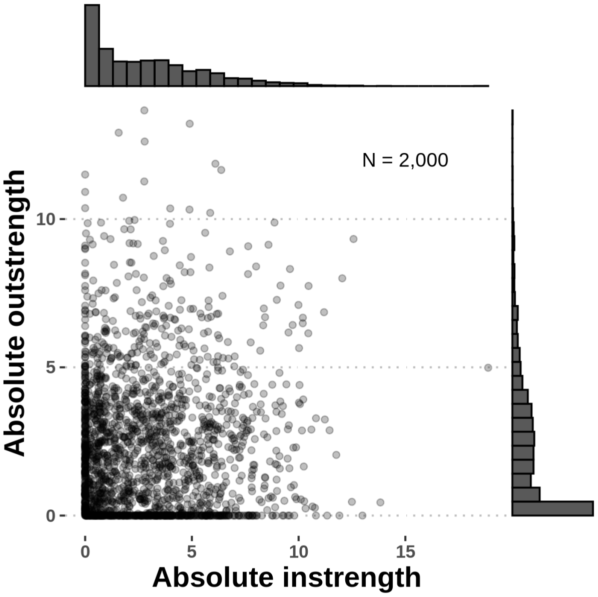


Figure S3: **Relationship between the node-level network metrics used in linear models.**
